# State-of-the-art estimation of protein model accuracy using AlphaFold

**DOI:** 10.1101/2022.03.11.484043

**Authors:** James P. Roney, Sergey Ovchinnikov

## Abstract

The problem of predicting a protein’s 3D structure from its primary amino acid sequence is a longstanding challenge in structural biology. Recently, approaches like AlphaFold have achieved remarkable performance on this task by combining deep learning techniques with coevolutionary data from multiple sequence alignments of related protein sequences. The use of coevolutionary information is critical to these models’ accuracy, and without it their predictive performance drops considerably. In living cells, however, the 3D structure of a protein is fully determined by its primary sequence and the biophysical laws that cause it to fold into a low-energy configuration. Thus, it should be possible to predict a protein’s structure from only its primary sequence by learning a highly-accurate biophysical energy function. We provide evidence that AlphaFold has learned such an energy function, and uses coevolution data to solve the global search problem of finding a low-energy conformation. We demonstrate that AlphaFold’s learned potential function can be used to rank the quality of candidate protein structures with state-of-the-art accuracy, without using any coevolution data. Finally, we explore several applications of this potential function, including the prediction of protein structures without MSAs.

---

Knowledge of 3D protein structures is critical for designing drugs, characterizing diseases, and creating a mechanistic understanding of cellular biology. Experimental approaches to protein structure determination can be costly and time-consuming, so the ability to computationally predict protein structures from amino acid sequences is extremely useful. Recently, AlphaFold demonstrated breakthrough performance on protein structure prediction, with predictions often nearing experimental accuracy [1]. Approaches like AlphaFold have advanced the state-of-the-art in protein structure prediction by using deep learning methods to analyze coevolutionary information. To predict the structure of a target amino acid sequence, these methods first search a database of protein sequences to compile a Multiple Sequence Alignment (MSA), which is essentially a collection of sequences that are evolutionarily related to the target sequence. MSAs are known to provide extremely useful information for predicting protein structures [2–4]. Intuitively, if two residues are in contact in a folded protein structure, mutations in the first position may induce a selective pressure for the second position to mutate. Such mutational covariance can be detected in MSAs, and this signal has been critical to the success of recent protein structure prediction models, including AlphaFold. However, the requirement of MSAs for protein structure prediction is sometimes problematic, since some proteins have few known homologs.

In theory, it should often be possible to predict protein structures without using MSAs, since protein structures are fully determined by their amino acid sequences [7]. More specifically, Anfinsen’s dogma states that protein structures fold to minimize free energy, which is a function of the protein’s 3D configuration and its amino acid sequence. Therefore, if one could model this energy function with sufficient accuracy, then one could predict protein structures by optimizing this function over the space of 3D configurations. Classical protein structure prediction methods like Rosetta take this approach, and sample structures from a hand-designed potential function [8]. The challenges with this approach are twofold. First, it is difficult to accurately model the biophysical energy function that governs protein folding at a level of abstraction that is computationally tractable. Second, even with perfect knowledge of the energy function, there are an astronomically large number of possible protein geometries, so searching for the optimum is a difficult global optimization task [9].

Given the theoretical possibility of predicting protein structures without MSAs, it is interesting to speculate why AlphaFold remains dependent on MSAs for its accuracy. One intriguing possibility is that AlphaFold has learned an accurate potential function for scoring the accuracy of candidate protein structures, but the coevolutionary information in the MSA is necessary to locate an approximate global minimum in this potential function and circumvent the challenging optimization problem. After finding the neighborhood of the global minimum using the MSA, the later stages of the AlphaFold model may act as an “unrolled optimizer” and locally descend the learned potential to produce a refined structure prediction. AlphaFold also outputs various confidence scores related to the predicted accuracy of its structures, and these confidence scores may be determined by the value of its internal potential function. This hypothetical prediction mechanism is illustrated in Figure 1.

**FIG. 1.**
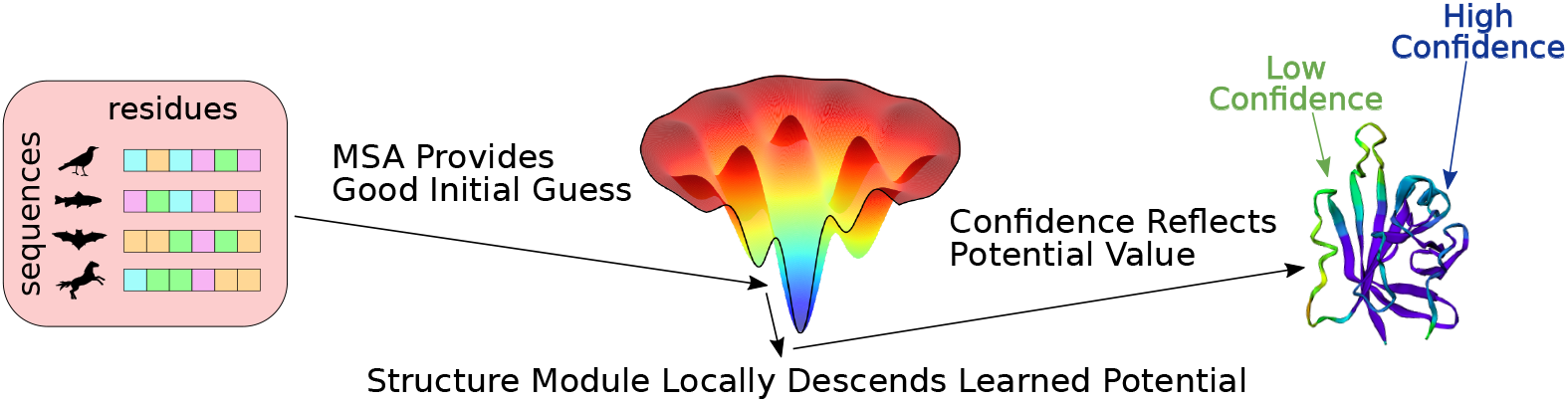
The hypothesized role of coevolutionary information in AlphaFold’s predictions. Images inspired by [5, 6].

This hypothesis lends itself to experimental testing. Candidate structures can be supplied to AlphaFold as templates, which are used to incorporate known structural information from proteins that are related to the target sequence. In this paper we show that, when a candidate structure is introduced as a template, AlphaFold’s confidence metrics are closely correlated with the actual accuracy of the candidate structure, even when no co-evolutionary information is supplied. This suggests that AlphaFold has learned an accurate biophysical potential function for protein structures that does not rely on coevolutionary information.

## Decoy Scoring

Computational biologists have historically predicted protein structures based on related sequences with experimentally solved structures [10]. AlphaFold incorporates this approach by allowing the structures of up to four related proteins to be supplied to the model as templates. For each template, AlphaFold receives the template’s one-hot-encoded amino acid sequence, C*β* distance matrix, and backbone and side chain torsion angles as inputs. In addition, AlphaFold is given a mask indicating which atoms are unresolved in the template structure, and ignores torsion angles involving those atoms. Recent papers have demonstrated that AlphaFold’s template mechanism can be used to refine experimentally and computationally derived structural hypotheses [11, 12].

We investigated whether AlphaFold has learned a coevolution-independent potential function for scoring protein structures by supplying AlphaFold with a.) a target amino acid sequence to be predicted and b.) a “decoy structure” that is passed to the model as a template. The goal of this procedure is to score the plausibility of the target amino acid sequence adopting the geometry given by the decoy structure. It is motivated by the hypothesis that AlphaFold’s output structure will resemble the decoy introduced as a template and therefore, if AlphaFold has learned an accurate potential function that does not require coevolution information, the output confidence metrics will closely track the quality of the decoy. Note that no coevolutionary information is supplied to the model during this procedure.

We used a sequence of all “gap” tokens (which represent missing amino acids) to fill in the one-hot-encoded amino acid sequence associated with the decoy. We used the gap sequence due to an initial observation that high sequence identity between the decoy sequence and the target sequence caused AlphaFold to be overconfident in the decoy’s accuracy (Figure S1). To keep the structural information supplied to AlphaFold from leaking the true decoy sequence, we masked out all side chain atoms aside from C*β*, and added a C*β* atom to all glycine residues (we decided to retain the C*β* atoms because AlphaFold uses a C*β* distance matrix to encode the template structure).

After processing its inputs, AlphaFold produces an output structure and two confidence metrics: the predicted TM Score (pTM) and the predicted LDDT-C*α* Score (pLDDT) [13, 14]. To determine whether AlphaFold has learned a MSA-free potential function for assessing protein structure accuracy, we investigated whether we could accurately rank the decoy structures based on AlphaFold’s outputs. For each decoy, we computed a “composite confidence score” by multiplying the output pLDDT, the output pTM, and the TM Score between the decoy structure and the AlphaFold output structure. The last term adjusts for the fact that AlphaFold’s confidence metrics ultimately reflect the accuracy of the output structure (which can differ from the decoy structure), while we were interested in scoring the decoy structures for the sake of direct comparison with other decoy-ranking methods.

### Rosetta Decoys

Using the procedure outlined above, we aimed to determine whether AlphaFold’s outputs could be used to assess the accuracy of decoy structures introduced as templates. For our initial evaluation we used the Rosetta decoy dataset, which contains 133 native protein structures with thousands of decoys for each native structure [15]. We compared AlphaFold’s ability to assess the quality of decoy structures with the Rosetta energy function, as well as DeepAccNet, which is a state-of-the-art machine learning model for estimating the accuracy of protein structure models [16]. All reported results are from AlphaFold model 1 with 1 recycling iteration.

We found the correlation between the composite confidence score and decoy quality to be robust and consistent. The average Spearman rank correlation between the composite confidence score and the quality of the decoy (as measured by TM Score to the native structure) was .925, compared to average correlations of .831 and .760 for DeepAccNet and the Rosetta energy function. Another practical indicator of decoy-ranking performance is the quality of the top-ranked decoy for each target. On the Rosetta decoy dataset, the top-ranked decoys selected via the composite AlphaFold confidence score had an average TM Score of .933 compared to .917 for DeepAccNet and .901 for Rosetta. More details on the Rosetta dataset are given in Figure 2.

**FIG. 2.**
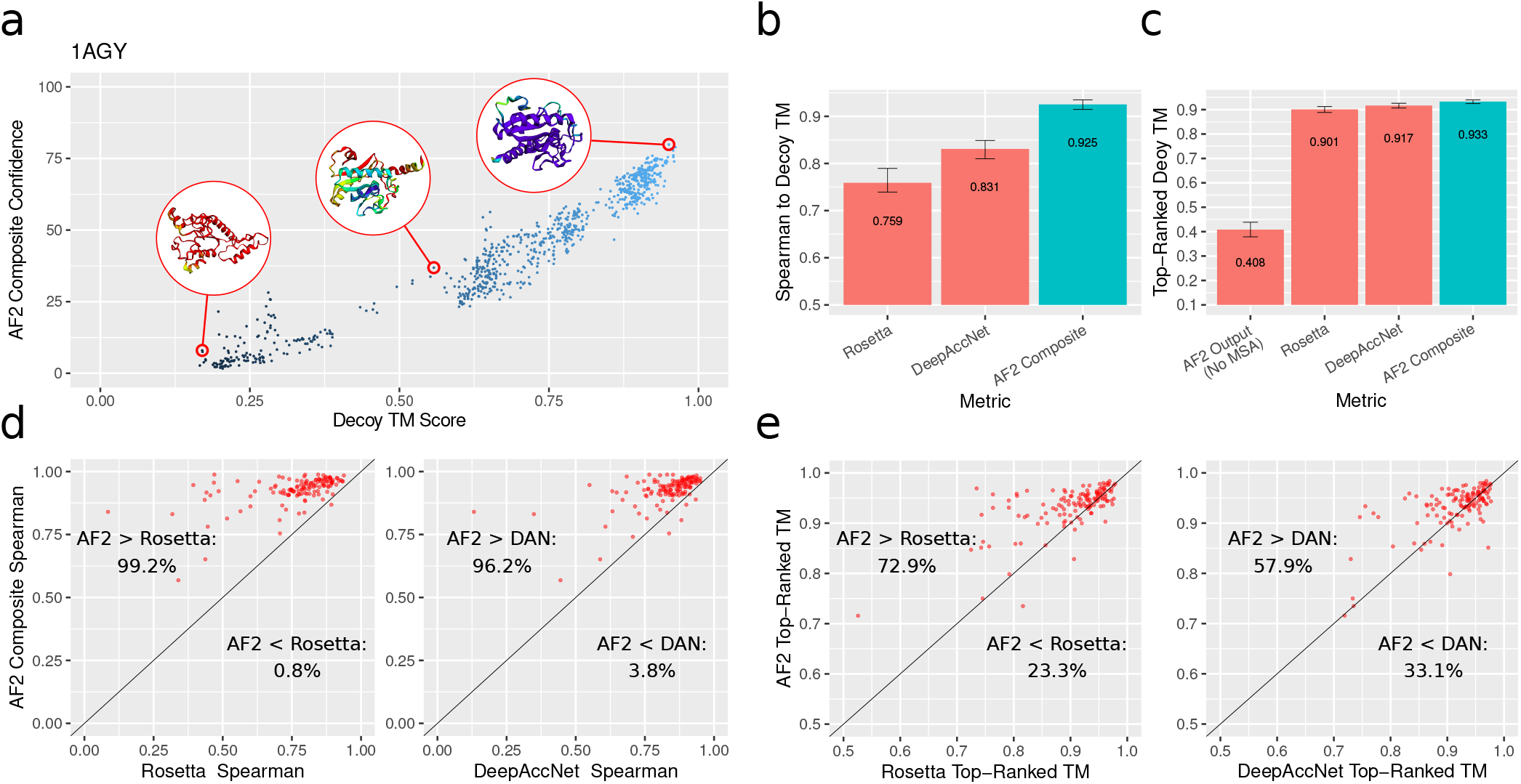
Decoy ranking results on the Rosetta decoy dataset. (a) Decoy TM Score vs. composite confidence for an example target. Three selected AlphaFold output structures are visualized, color indicates model confidence. (b) Mean Spearman correlations between various metrics and decoy TM Score. (c) Mean TM Scores of the top-ranked decoys for various metrics, as well as the mean TM Score of AlphaFold’s prediction with no MSA. (d) Comparison of Spearman correlations for AlphaFold and Rosetta/DeepAccNet. (e) Comparison of top-1 accuracies for AlphaFold and Rosetta/DeepAccNet. All error bars are bootstrap 95% confidence intervals of the mean.

Overall, these evaluations indicate that AlphaFold can assess the quality of candidate protein structures with state-of-the-art accuracy, even when no coevolution information is provided. It should be noted that AlphaFold’s structure predictions were of low quality when no templates were provided (average TM Score of .408). Yet despite being unable to predict the structures of these proteins without an MSA, AlphaFold achieved excellent performance assessing the quality of decoys without any MSA inputs. This provides evidence for the hypothesis that AlphaFold has learned a potential function that is largely independent of coevolution information, but needs coevolution information to search for global optima in this potential.

### CASP1

To assess the decoy-ranking ability of AlphaFold on a novel sample of proteins, we performed an additional evaluation on the Estimation of Model Accuracy (EMA) task from CASP14 [17]. To set up the CASP14 EMA experiment, the CASP organizers created a set of decoy structures by taking the 150 most accurate server submissions for each structure prediction target in CASP14. Note that the decoy set does not include predictions from AlphaFold, since AlphaFold was entered in CASP14 as a human group rather than a server. We replicated this evaluation using AlphaFold (with the gap sequence) to assess the decoy structures, and compared the results with ranking methods entered in CASP14.

The CASP assessors evaluated EMA methods based on their top-1 GDT TS loss, which is the difference in GDT TS scores between the best decoy and the topranked decoy by a given EMA method [18]. EMA methods were ranked based on their average GDT TS loss over targets where at least one decoy had GDT TS over 0.4, as well as the average Z-Score of their GDT TS loss over these targets. For both metrics, the AlphaFold composite confidence score significantly outperformed all other EMA methods entered in CASP14. Results from the CASP14 evaluation are presented in Figure 3.

**FIG. 3.**
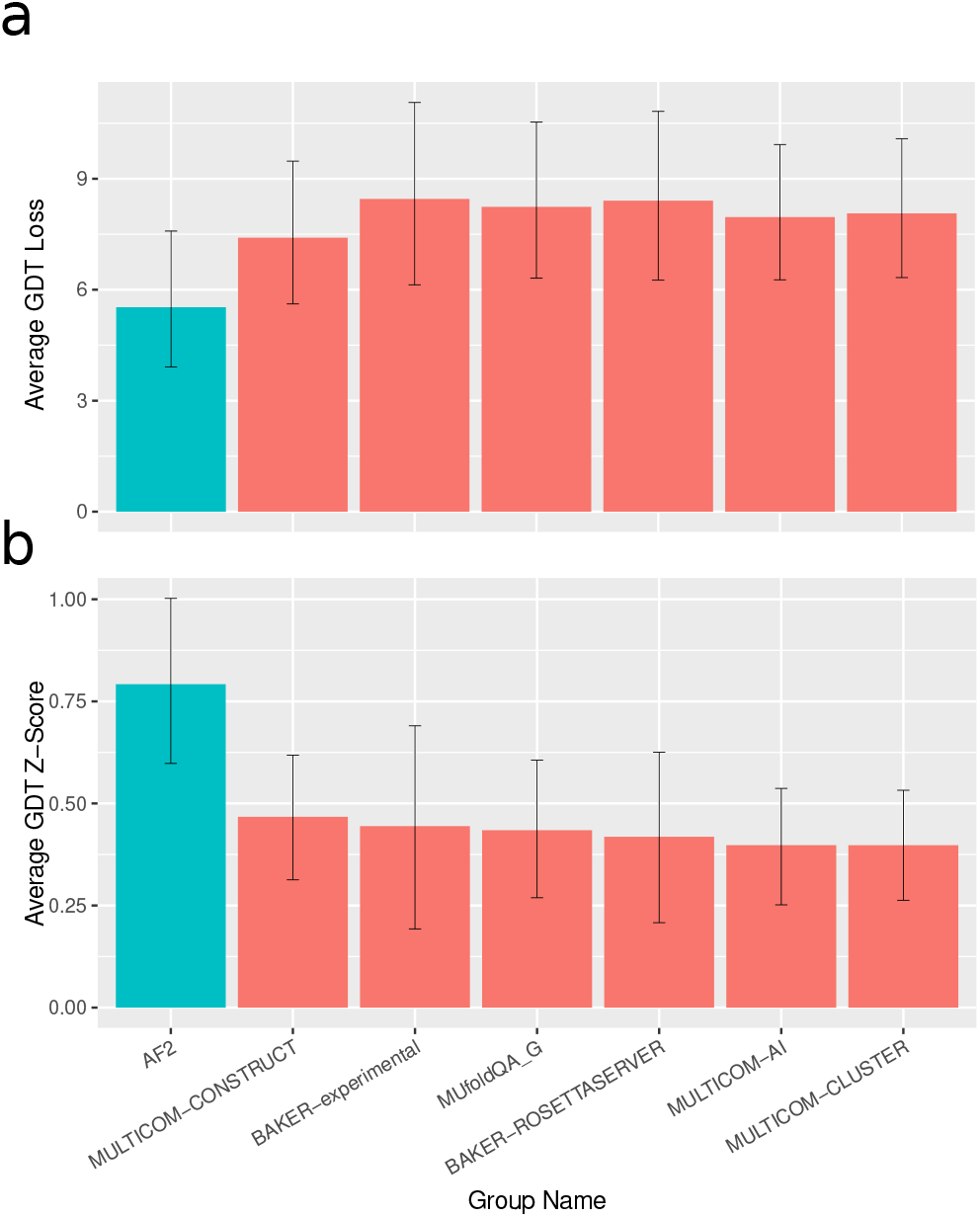
Decoy ranking results on CASP. (a) GDT TS loss for AlphaFold and top EMA methods from CASP14. (b) GDT TS Z-Scores for AlphaFold and top EMA methods from CASP14. Error bars are bootstrap 95% confidence intervals of the mean.

These results indicate that AlphaFold can reliably assess the accuracy of candidate protein structures without the use of coevolution information. However, coevolution data (or a method that can generate decoys close to the correct structure) are still necessary for accurate structure prediction, since AlphaFold generally failed to predict accurate structures for the CASP14 targets without an MSA (Figure S3).

### Applications

Our finding that AlphaFold can assess the accuracy of candidate protein structures without the need for coevolution data opens up several exciting applications. One such application is the prediction of protein structures without MSAs. In theory, it should be possible to accurately predict protein structures by searching over the space of possible decoy structures and finding those that are highest-ranked by AlphaFold. However, given the vast number of possible candidate structures, an exhaustive search is intractable.

One way of mitigating this intractability is to search over the output space of a generative model of realistic protein structures. Instead of training a new generative model of candidate structures, we designed a generator-discriminator pipeline that links two instances of AlphaFold (Figure 4B). The first instance of AlphaFold (the generator) takes an arbitrary amino acid sequence as input, and produces a candidate protein structure as output. This candidate structure is then supplied to the discriminator as a template (with a sequence of gap tokens). Finally, the discriminator tries to predict the structure of the target sequence using the template, and produces confidence outputs in the process. As demonstrated by our previous experiments, these confidence metrics are strongly correlated with the accuracy of the candidate structure. By perturbing the input sequence to the generator, we can explore the space of candidate structures while using the discriminator’s confidence metrics as an indicator of accuracy. We performed this exploration by backpropagating the discriminator’s confidence signal to the input sequence and updating it via stochastic gradient ascent, thereby molding the input sequence to produce a high-quality candidate structure from the generator.

**FIG. 4.**
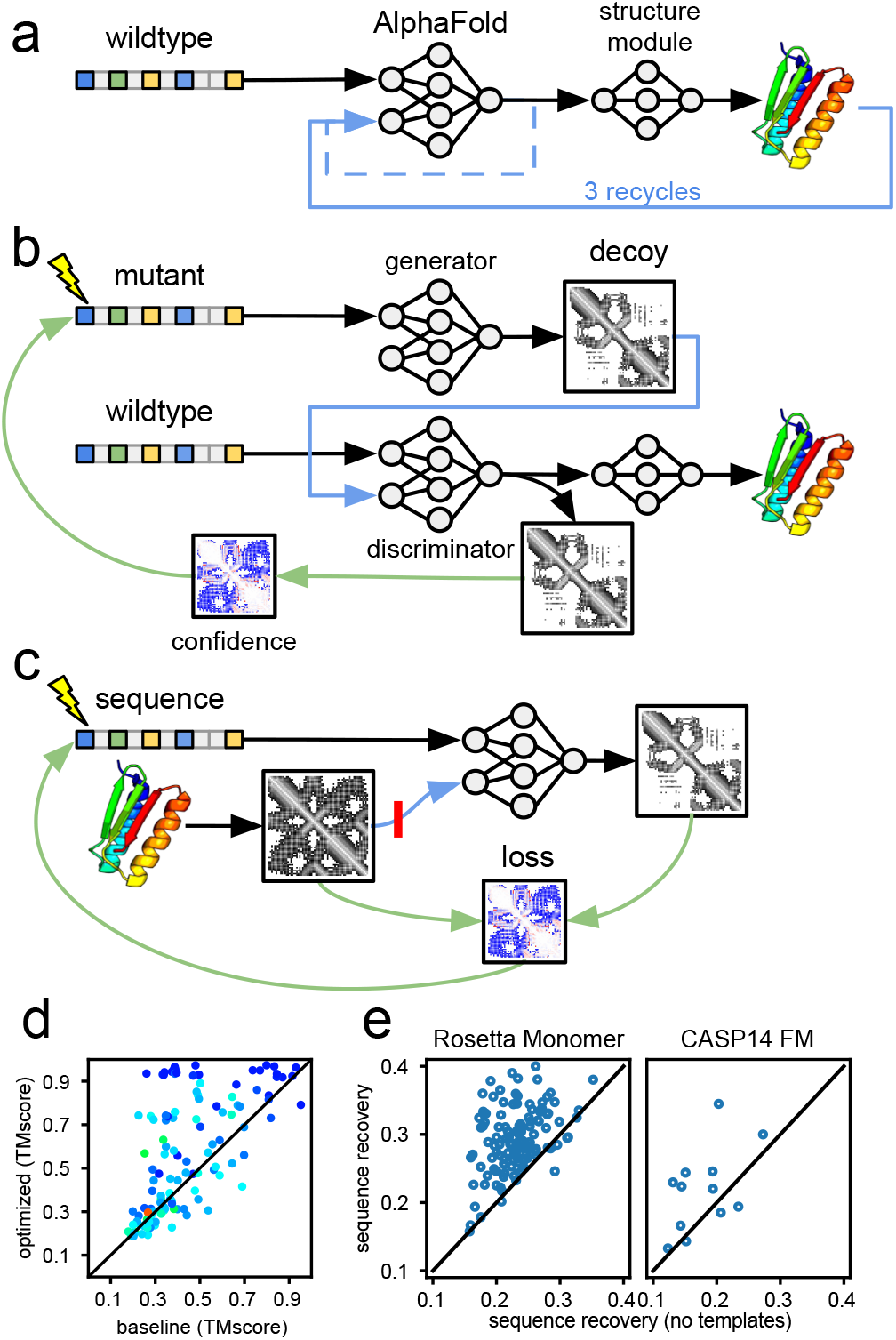
Application of AlphaFold’s template mechanism for sequence and structure generation. Single-sequence prediction with (a) the baseline structure prediction protocol with 3 recycles or (b) two instances of AlphaFold for structure generation and discrimination. (c) Protocol for sequence design to minimize loss between desired and predicted structure via distogram, with and without template (red line). (d) Comparing structure accuracy of (a) vs (b) on the Rosetta decoy set. Dots colored by pLDDT red to blue (50 to 90). (e) Comparing sequence recovery with and without templates on the Rosetta monomeric and CASP14 FM datasets.

We found that, with this approach, we were able to improve upon AlphaFold’s structure predictions when no MSAs were available. Although we did not use recycling in the generator and discriminator models, we compared the quality of our optimized predictions to a baseline created by running AlphaFold with a single sequence input and 3 recycling iterations. On the Rosetta decoy set, we could significantly improve prediction quality (ΔTMScore *>* 0.1) on 50 examples out of 123 compared to running the baseline (Figure 4D). We hypothesize that this procedure is able to improve AlphaFold’s predictions because it performs a more wide-ranging search than the “unrolled optimizer” implemented by AlphaFold itself. Though our results demonstrate the potential of searching over candidate structures using AlphaFold’s learned potential function, the current optimization protocol sometimes gets stuck in local minima with low accuracies and low confidence scores (Figure S9-11).

Since AlphaFold’s learned potential function can determine the level of compatibility between a protein structure and sequence without the need for coevolution data, it is also applicable to protein design (i.e., the problem of finding a protein sequence that folds into a target backbone geometry). Our work suggests a straightforward approach to protein design using AlphaFold: supply the desired backbone structure to AlphaFold as a template (with a sequence of gap tokens and side chains masked), and optimize the composite confidence score with respect to the input sequence. To facilitate gradient-based optimization, we used the categorical cross-entropy between AlphaFold’s predicted distance matrix and the template distance matrix as a differentiable surrogate for the composite confidence score. This design procedure achieved substantial levels of native sequence recovery in the neighborhood of 30%.

Repeating the same procedure without a template input resulted in significantly lower sequence recovery (Figure 4E). This is likely because, without a template or MSA, AlphaFold often fails to predict the correct structure for the input sequence. This leads to “false negatives” while optimizing the input sequence to match the target backbone (i.e., input sequences that would actually fold into the target backbone are mis-predicted by AlphaFold and incorrectly assigned high loss). The template input eliminates false negatives by providing a good starting point for AlphaFold’s structural optimization, allowing AlphaFold to confidently and accurately predict when an input sequence will fold into the target structure. The effectiveness of template inputs at increasing sequence recovery supports our hypothesis that AlphaFold has learned a potential function that can assess sequence-structure agreement, but needs coevolution data or templates to help search for optimal structures.

## Conclusions

We have demonstrated that AlphaFold has learned a protein structure potential that does not need coevolution information to achieve high accuracy, although AlphaFold still needs coevolution data to search for global minima in this potential. This finding has significance for the interpretation of protein structure prediction models, as well as practical applications. These applications include the prediction of protein structures when MSAs are not available and the improvement of protein design methods.

## Code Availability

The code used to run the evaluations in the paper, as well as the raw data, is available at https://github.com/jproney/AF2Rank.

## Revision Summary

The current version of this draft has some noteworthy differences from the previous version. In our original draft, we investigated two choices for the one-hot encoded sequence associated with the decoy structures: a sequence of all alanines, and the target sequence. We later discovered that we had used an incorrect encoding for the target sequence (this error is described in detail in the accompanying code release). Consequently, our reported results for the target sequence are significantly different in this draft. We also switched from using a sequence of all alanines to a sequence of all gap tokens to avoid any bias towards alanine-rich sequences, although these sequences gave highly similar results. Our previous draft also experimented with combining multiple choices of decoy sequence to create even stronger ranking results, but we ultimately decided it was simpler to focus on the results from using the gap sequence. Finally, we added an applications section to the main text, which expands upon the decoy generation results that were previously included in the appendix. This revision also includes new experiments exploring applications to protein design and mutation effect prediction, the latter of which is described in Appendix E.

## Acknowledgement

We would like to thank John Jumper for helpful comments on our original manuscript. SO is supported by NIH Grant DP5OD026389, NSF Grant MCB2032259 and the Moore–Simons Project on the Origin of the Eukaryotic Cell, Simons Foundation 735929LPI, https://doi.org/10.46714/735929LPI.

## Appendix A: Comparison of Decoy Sequences

As mentioned in the main text, we investigated two choices for the one-hot encoded sequence associated with the decoy structure: the target amino acid sequence, and a sequence of “gap” tokens. While AlphaFold’s confidence metrics were robustly correlated with decoy quality when using the gap sequence, this correlation was much lower when using the target sequence. A comparison of decoy-ranking performance for each of the two sequences on an example protein is given is Figure S1.

**FIG. S1.**
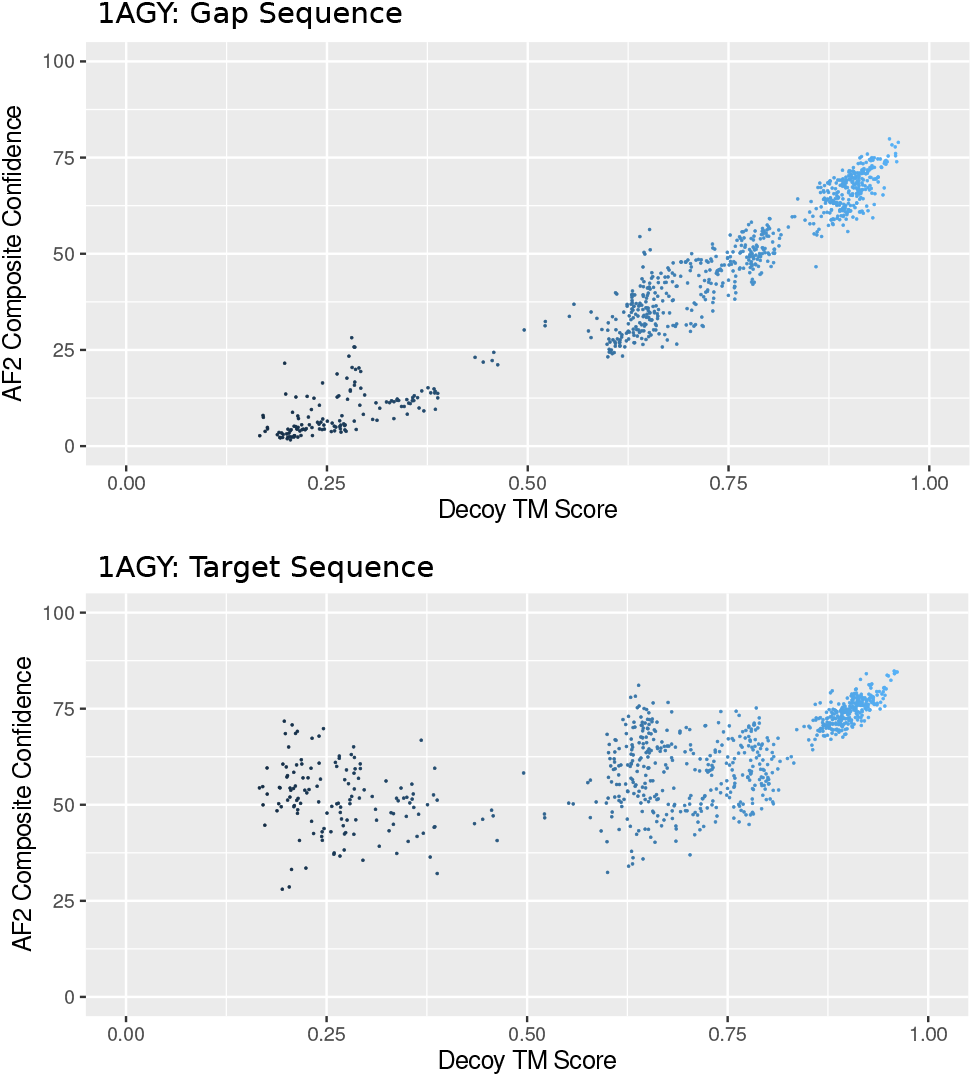
Correlation between AlphaFold’s confidence metrics and decoy quality when using the gap sequence (top) and the target sequence (bottom).

We hypothesize that AlphaFold becomes overconfident in the quality of the decoy when there is high sequence identity between the decoy and the target. Due to this difference in performance, we used the gap sequence in our evaluations.

## Appendix B: Analysis of Output Structures

AlphaFold’s output structures can differ from the structures provided as templates. In our experiments, AlphaFold’s output structures were often similar in quality to the decoy structures, and sometimes were substantially improved in terms of TM Score and GDT TS (Figure S2). While AlphaFold was sometimes capable of improving decoy structures without coevolution information, it generally could not predict these structures from scratch when no coevolution information was provided. This supports the idea that AlphaFold can perform local optimization over its learned protein potential, but needs coevolution data or a template to locate a good starting point for this optimization.

**FIG. S2.**
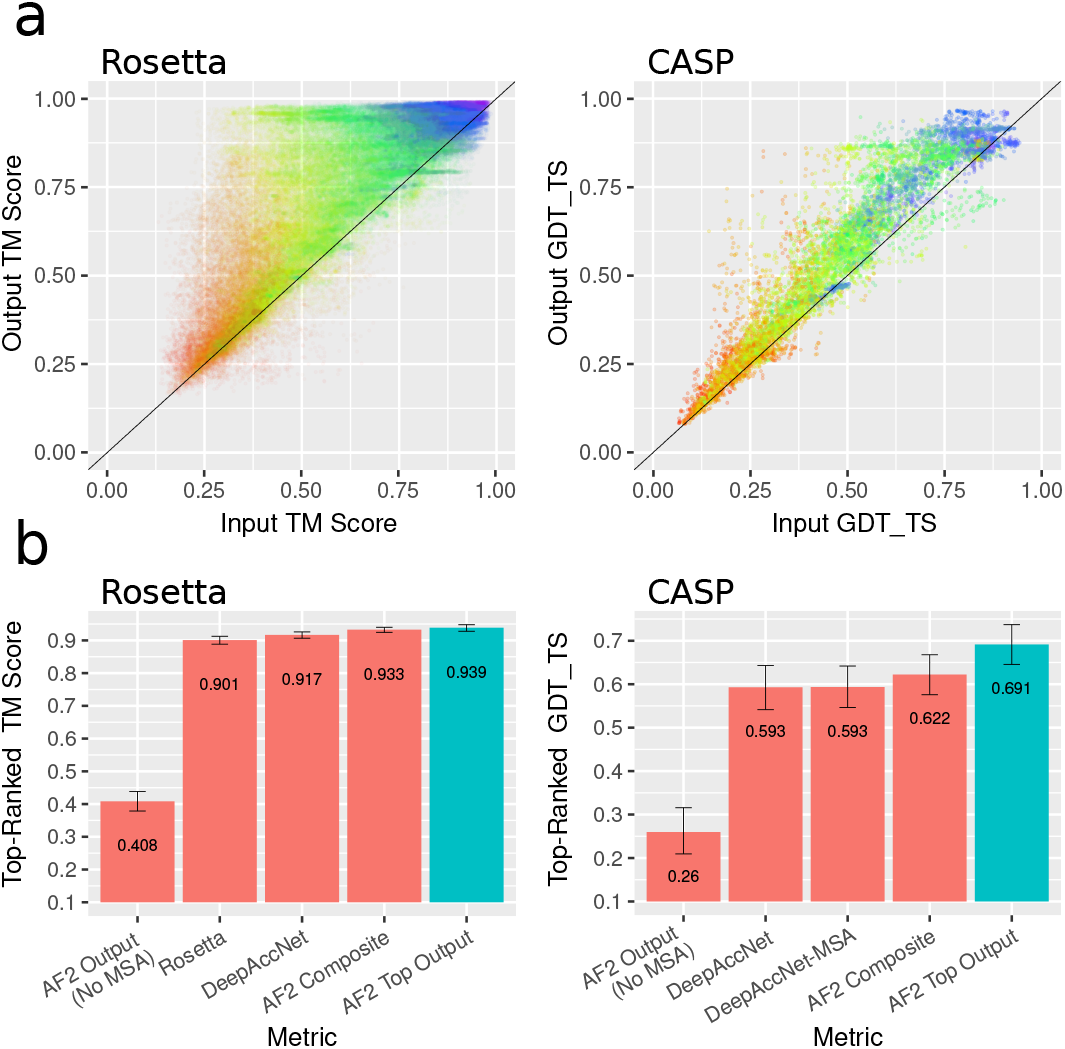
Comparison between input and output structure qualities. (a) TM Score / GTS TS of AlphaFold output structure vs. TM Score / GDT TS of decoy structure supplied as a template. Each dot is single decoy in the Rosetta decoy set (left) or CASP EMA set (right), color indicates composite confidence score. (b) Mean TM Scores of the top-ranked Rosetta (left) and CASP (right) decoys for various ranking metrics, including the AlphaFold output structures with the highest pLDDT *×* pTM product in blue. Error bars are bootstrap 95% confidence intervals of the mean.

Although AlphaFold’s confidence metrics track the quality of its output structures, our composite confidence score effectively tracks the quality of the *decoy* structures (i.e., the color gradient in Figure S2A progresses along the x-axis). This is due to the inclusion of a term that measures the change between the decoy structure and the output structure.

## Appendix C: CASP Evaluation Extended Results

In this section we provide more details on the results of our CASP14 evaluation. As described in the main text, we found that AlphaFold outperformed all methods from the CASP14 EMA experiment (according to the metrics used by CASP). We also directly compared AlphaFold to DeepAccNet (entered in CASP14 as BAKER-ROSETTASERVER) and DeepAccNet-MSA (entered as BAKER-Experimental), which were two of the top-performing methods in the CASP14 EMA experiment. As shown in Figure S3, AlphaFold outperformed both of these methods on a majority of targets.

**FIG. S3.**
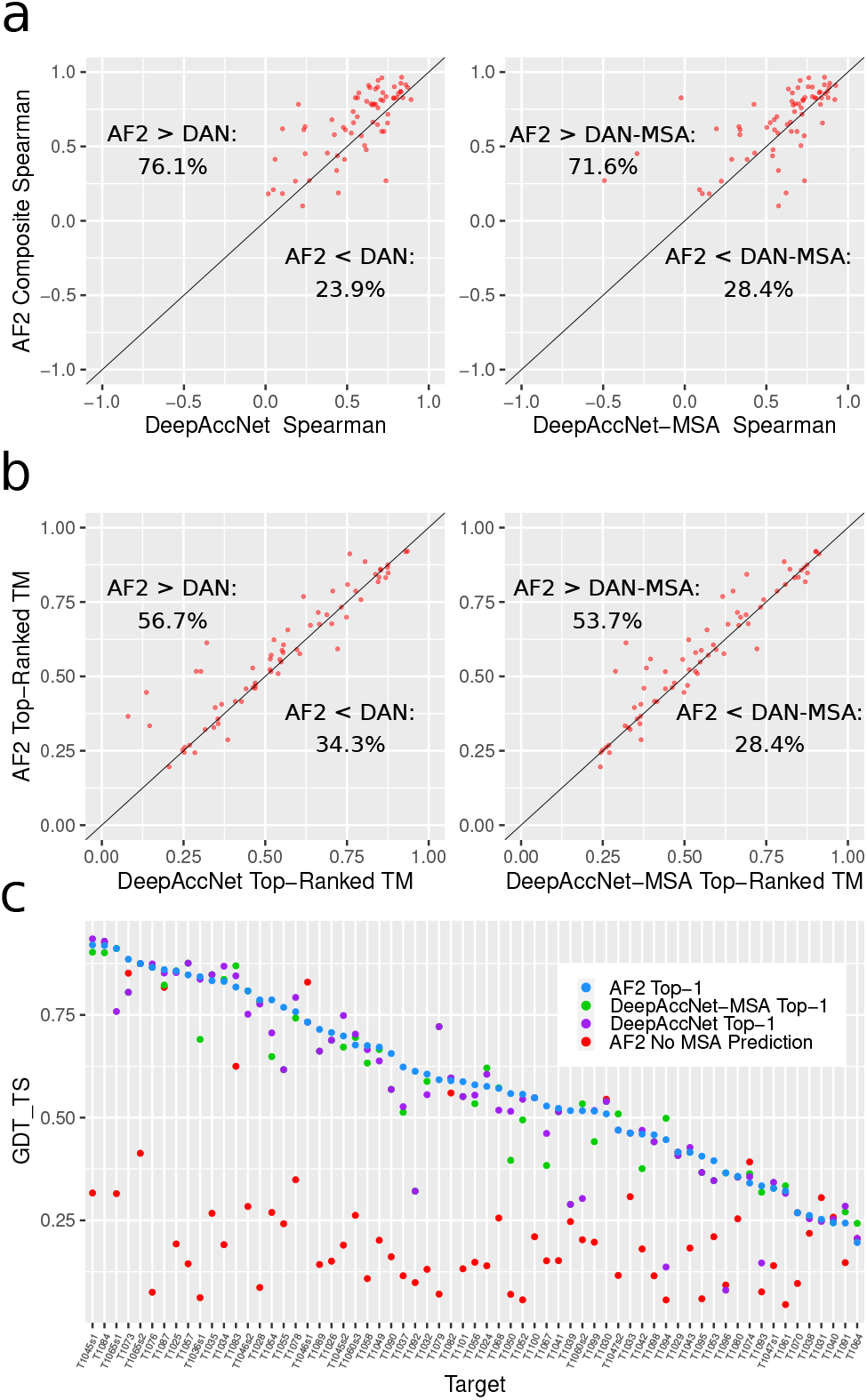
Extended Results from the CASP14 Evalutation. (a) Comparison of Spearman correlations for AlphaFold and DeepAccNet/DeepAccNet-MSA. (b) Comparison of top-1 accuracies for AlphaFold and DeepAccNet/DeepAccNet-MSA. (c) Top-1 results on each CASP14 target for AlphaFold, DeepAccNet, and DeepAccNet-MSA, as well as the MSA-free AlphaFold prediction for each target.

These results show that AlphaFold can reliably assess the accuracy of candidate protein structures without the use of coevolution information. However, coevolution data (or a method that can generate decoys close to the correct structure) are still necessary for accurate structure prediction. When AlphaFold was tasked with predicting the CASP14 targets without any MSA inputs, its structure predictions were generally much less accurate than the top-ranked decoys based on AlphaFold’s confidence metrics (Figure S3C).

In the main text, we used the AlphaFold composite confidence score with a “gap” decoy sequence to rank the CASP decoys. We chose this configuration because it gave the best performance on the Rosetta dataset, which we used for validation. For completeness, Figure S4 shows the performance of other variations of the AlphaFold ranking scheme on the CASP dataset. In particular, we also tried using a sequence of all alanine residues, as well as using only the pLDDT and pTM scores in the composite confidence score. All of these configurations gave state-of-the-art results.

**FIG. S4.**
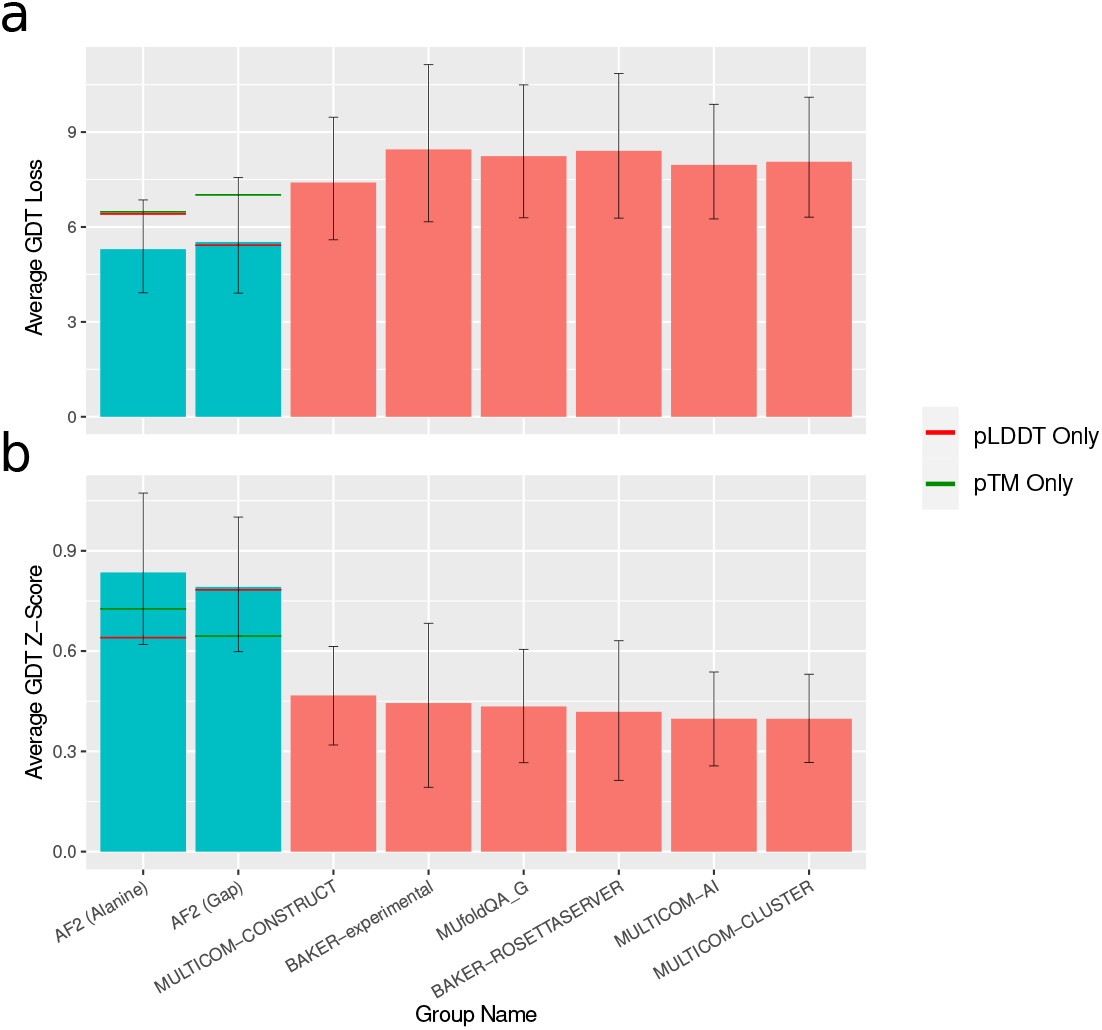
CASP14 performance for several variations of the AlphaFold ranking system. (a) GDT TS Loss (b) GDT TS Z-Scores.

## Appendix D: Additional Rosetta Ranking Results

**FIG. S5.**
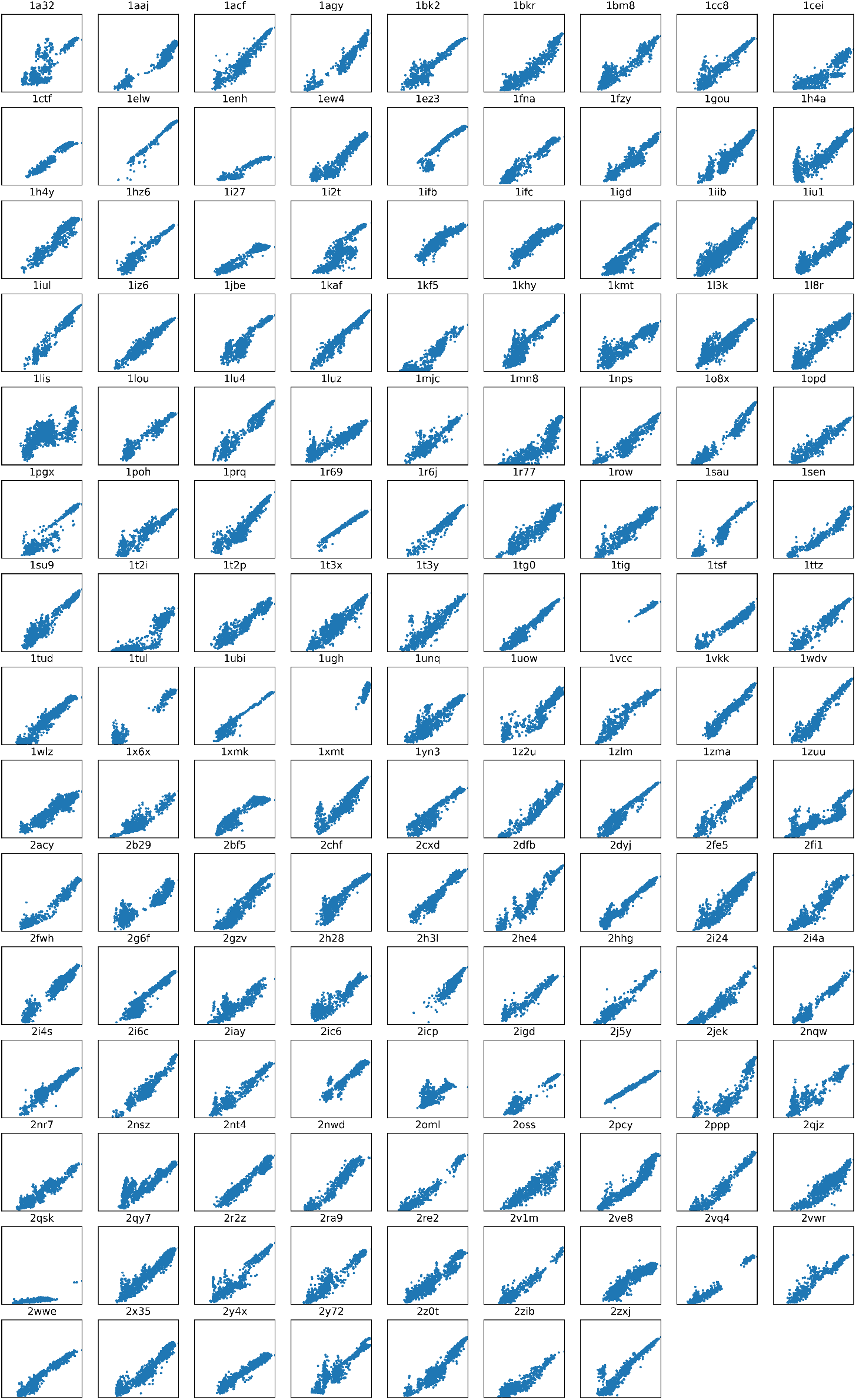
AF2 composite confidence vs decoy TM Score for all targets in the Rosetta decoy set.

## Appendix E: Mutant Effect Prediction

Our results provide evidence that AlphaFold has learned a potential function somewhat akin to a thermodynamic free energy, which is capable of assessing the accuracy of candidate protein structures without the need for coevolution information. A natural question is whether this “energy function” is capable of predicting the effects of point mutations on protein structure and stability. To explore this question, we attempted to use AlphaFold’s confidence metrics to predict the effects of point mutations on the fitness of *β*-Lactamase. We used the dataset from [19], which contains fitness assays for all point mutations to *β*-Lactamase.

**FIG. S6.**
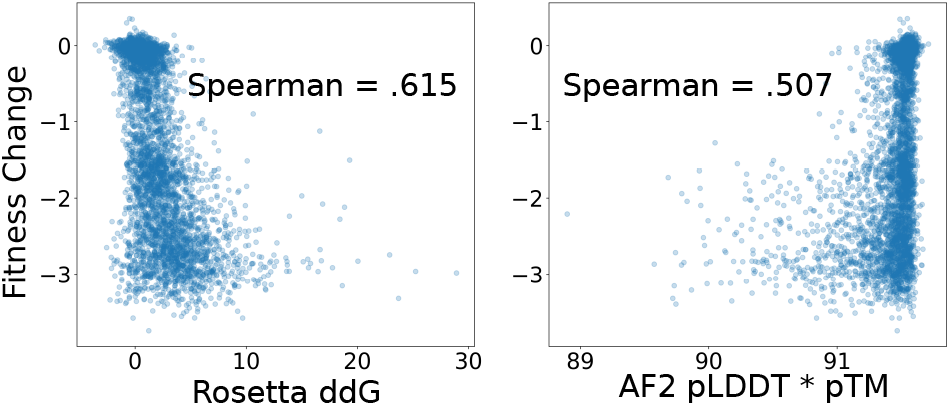
Correlation between *β*-Lactamase mutant fitness and Rosetta ΔΔG (left) and AF2 confidence metrics (right).

To predict the fitness effect of a given point mutation, we supplied AlphaFold with the native structure of *β-*Lactamase as a template (setting the template sequence to the native sequence and masking the side chains). For each mutation, we set the target sequence to the mutated sequence and recorded the output pLDDT and pTM. Based on our previous results, we might expect that the introduction of a destabilizing mutation into the target sequence will make it less compatible with the native structure supplied as a template, resulting in a drop in confidence scores. We ranked the mutant sequences by their pLDDT *×* pTM products, and computed the rank-order correlation of these scores with mutant fitness. We used the Rosetta ΔΔG values computed by [20] as physics-based baseline for mutation effect prediction.

We found that the pLDDT *×* pTM product of AlphaFold’s confidence metrics achieved a moderate correlation with fitness measurements of 0.51. The Rosetta ΔΔG values had a somewhat higher correlation of 0.62. This result suggests that, while AlphaFold’s learned potential function is more accurate than Rosetta at predicting the global accuracy of protein structures, Rosetta is somewhat better at predicting the impacts of point mutations. This is not altogether unsurprising, since AlphaFold was trained on stable structures, so its learned potential may be less sensitive to the effects of single destabilizing mutations and better suited to analyzing more global features of the sequence and structure.

## Appendix F: Protein Design Experiments

As described in the main text, we applied AlphaFold’s learned potential function to design protein sequences that fold into a set of target backbone structures. For each target structure, AfDesign was used to generate sequences that AlphaFold predicts will fold into the desired structure (available at https://github.com/sokrypton/ColabDesign/tree/beta/af). For this task, we used the fixbb (fixed backbone) protocol and the 3-stage design method. This protocol uses a categorical cross-entropy loss between the desired and the predicted distogram (a distogram is a tensor that contains a binned distribution of distances for every pair of residues). The protocol optimizes this loss over an input sequence (encoded as a *N ×* 20 matrix) that is used as input for both the target features and MSA features of AlphaFold.

During the 3-stage design protocol, the sequence-matrix is initially an unconstrained and continuous set of logits. Over time, the sequence-matrix is gradually constrained to become a normalized probability distribution according to the formula

~~~
(1-p) * logits + p * softmax(logit/temp).
~~~

For the first 300 iterations p is linearly scaled from 0 to 1, resulting in a softmax distribution at stage 2. For the next 200 iterations the temperature is reduced from 1.0 to 0.01, so the sequence-matrix approaches a one-hot encoded sequence. At the third stage, the one-hot encoded sequence is directly optimized for 50 steps, using the straight-through trick. At each step of optimization (across all 3 stages), only one AlphaFold model is used, but the model parameters are randomly selected from either model 1 or 2 (since these are the only ones trained with template inputs). To help the optimizer escape local minima, dropouts are enabled throughout the model. When a template input is used, 15% of the sites are randomly dropped at each iteration. At the third stage, the dropouts are disabled and the sequence with best loss is selected as the final design. Using 5 random seeds, this design protocol was repeated 5 times, with and without template inputs, to generate a total of 10 sequences.

**FIG. S7.**
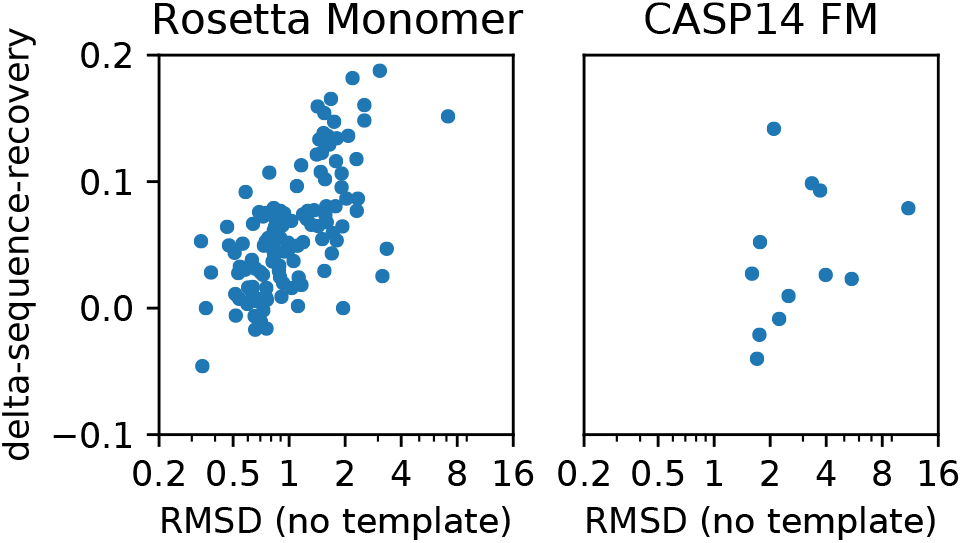
In cases where sequence optimization was relatively unsuccessful (i.e., the designed sequence still had high predicted RMSD with the target backbone), supplying the target backbone as a template improved sequence recovery.

As depicted in Figure 4E, we found that utilizing a template resulted in substantially higher levels of native sequence recovery. This improvement was especially large in cases where template-free optimization failed to find an input sequence that folded closely into the target backbone geometry (Figure S7).

Sequences were generated for both the proteins in the Rosetta decoy and CASP14 datasets. For CASP14, domains labelled as FM (free-modeling) were selected, as these are significantly different from any proteins in the AlphaFold training set. To reduce runtime, only proteins/domains of length 150 or less were used.

## Appendix G: MSA-free Structure Prediction

To improve AlphaFold’s structure predictions without the need for MSAs, we designed a generator-discriminator pipeline that links two instances of AlphaFold. By iteratively perturbing the input sequence-matrix, the generator is used to sample and refine decoy structures. Each generator output structure is passed to the discriminator as a template (with side chain atoms beyond C*β* masked and a sequence of gap characters). The discriminator then attempts to predict the target sequence, and its confidence is backpropagated to update the input sequence. Before backpropagation, the input sequence is initialized to the target sequence.

To make backpropagation easier, we utilized a confidence loss based on the discriminator’s distogram prediction head. The entropies of AlphaFold’s distance predictions convey model confidence, and using this signal as a loss does not require backpropagating through the structure module. More specifically, the loss function calculates the discriminator confidence for a given position by outputting the lowest entropy of any predicted distance distribution between that position and another residue more than 9 positions away. This entropy is calculated over the subset of bins in the distogram corresponding to distances less than 14 angstroms. Thus, the confidence loss is defined as:

~~~
loss = -(softmax(logits[bins<14])*
log(softmax(logits))[bins<14]).
~~~

This formulation assesses AlphaFold’s confidence in its predictions of tertiary contacts.

We experimented with two different ways of linking the instances of AlphaFold together: 1) by passing the predicted distogram from the generator to the discriminator model as a template distogram, and 2) by passing the predicted distogram and pair representations from the generator into the discriminator via AlphaFold’s recycling mechanism. Both approaches avoid backpropagating through the structure module by directly passing the predicted distogram, making optimization smoother.

For the input sequence optimization, we experimented with two approaches: 1) greedily optimizing random point mutations and 2) backpropagating the loss and updating the input sequence using stochastic gradient descent. For the first approach, at each iteration, 10 random mutations were evaluated and the mutation resulting in the minimal loss was fixed. 10 independent trajectories with 50 iterations each were performed. The discriminator output structure with the best loss across all iterations and trajectories was selected (Figure S8A, Figure S9). For the second approach, the input sequence-matrix was treated as an unconstrained set of continuous values during optimization. 50 independent trajectories were carried out for 50 steps each with a learning rate of 0.1 (Figure S8B, Figure S10). The same was performed for the recycle experiment (Figure S8C, Figure S11).

**FIG. S8.**
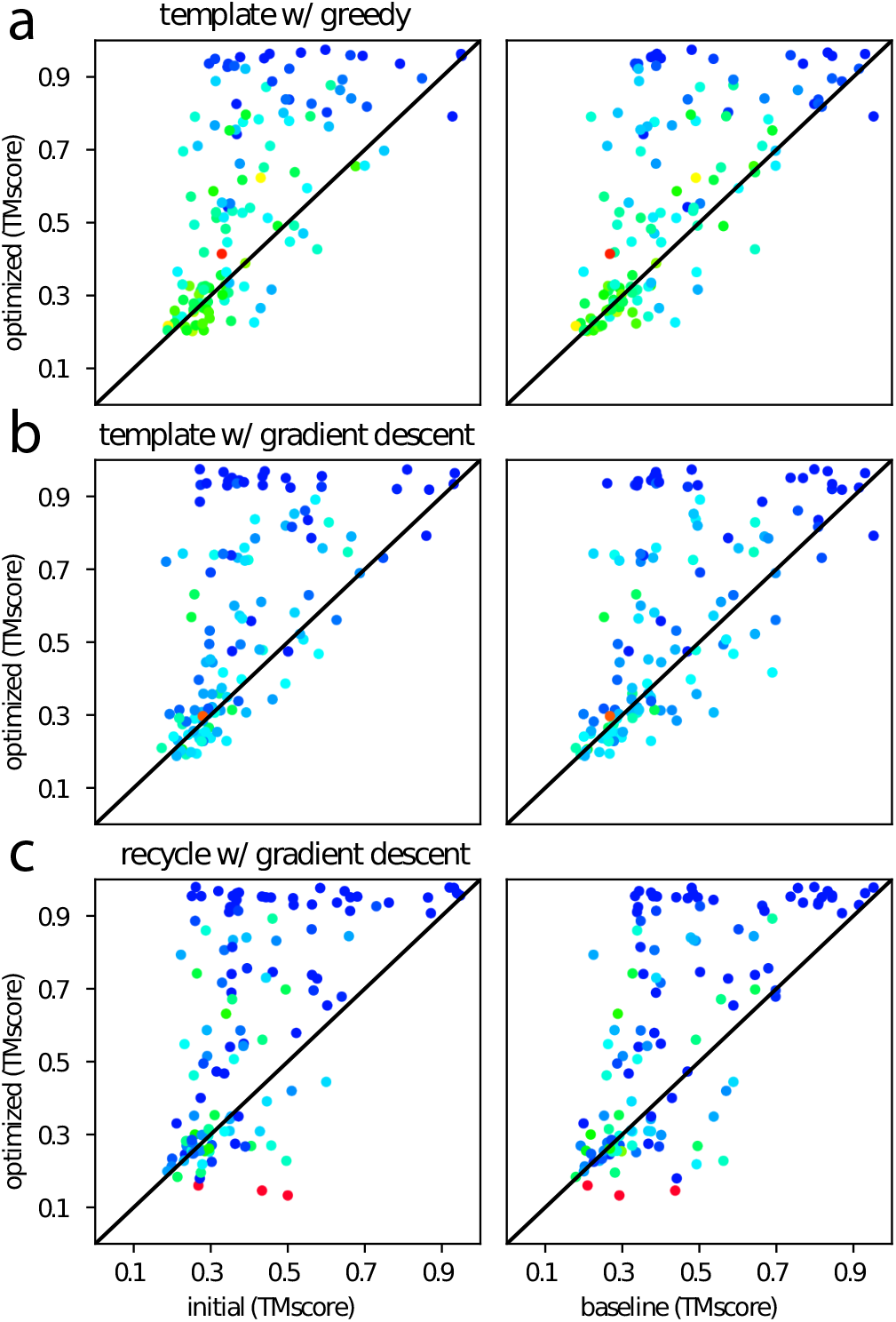
Decoy generation using AlphaFold improves structure predictions from single sequences, regardless of sampling techniques. (a) Semi-greedy mutant optimization. (b) Gradient descent via the template mechanism. (c) Gradient descent via the recycling mechanism. Left column compares structure accuracy before and after optimization. Each dot is one of the 133 proteins in the Rosetta decoy set. To control for the fact that linking two models is similar to an instance of recycling, the right column compares the predicted structure after optimization to the prediction from the baseline AlphaFold protocol with 3 recycles and a single-sequence input. The color is the discriminator pLDDT (red to blue, 40 to 90)

**FIG. S9.**
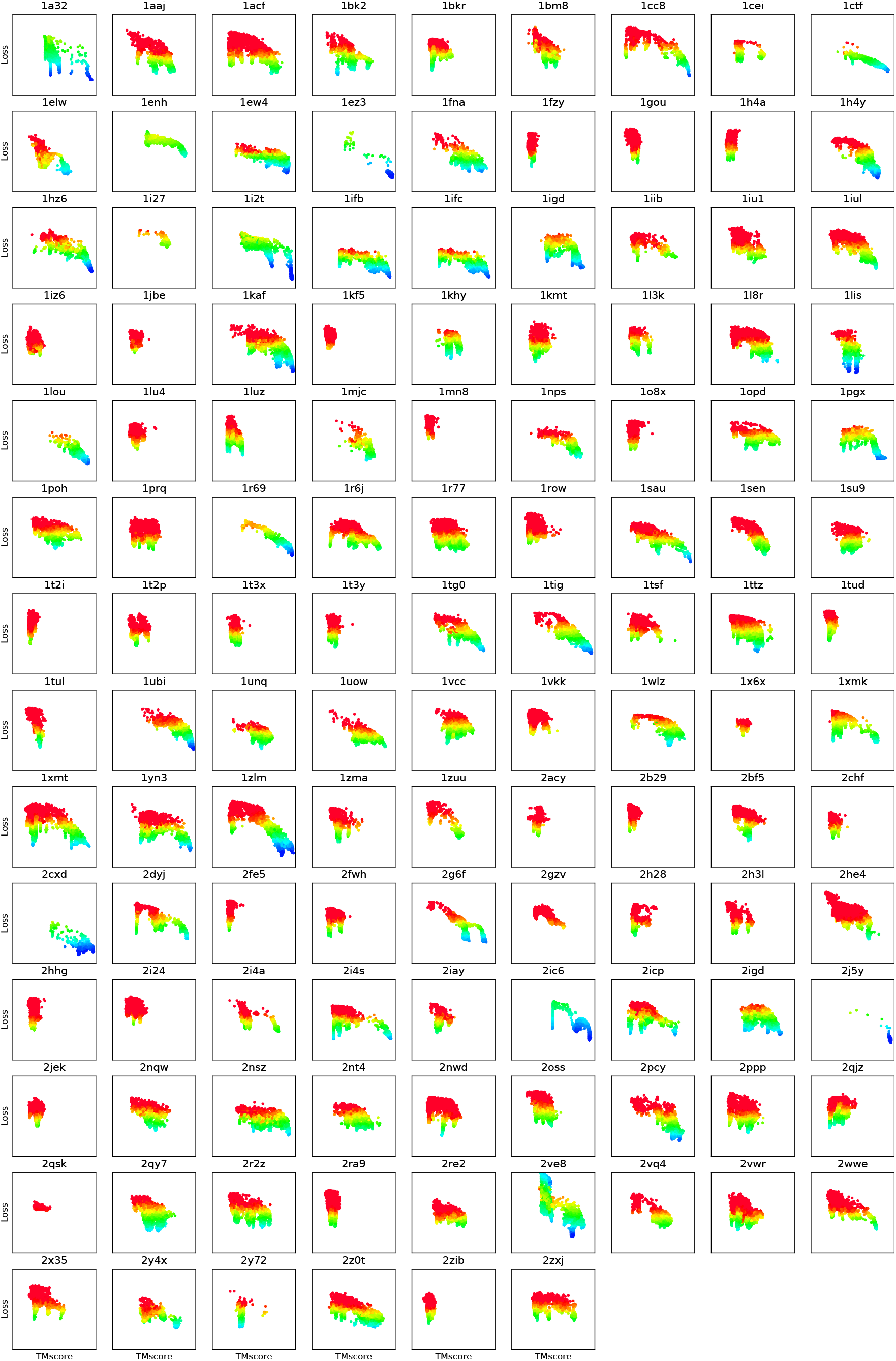
Ranking AlphaFold generated decoys with semi-greedy mutant optimization via templating mechanism. X-axis is the TMscore (range 0 to 1), y-axis is loss (range 1.4 to 4.6), color is the discriminator pLDDT (red to blue, 40 to 90)

**FIG. S10.**
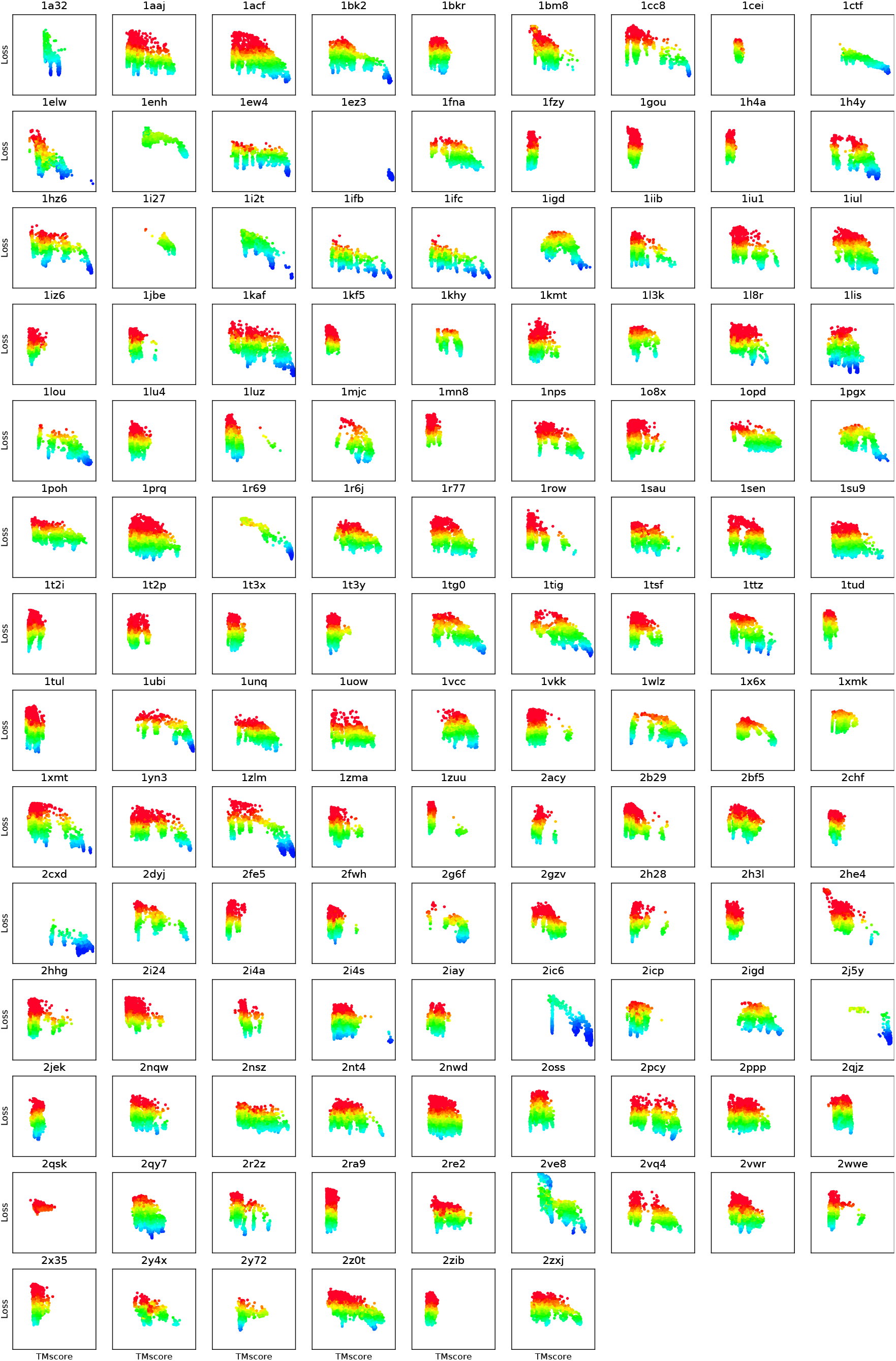
Ranking AlphaFold generated decoys with gradient descent optimization via templating mechanism. X-axis is the TMscore (range 0 to 1), y-axis is loss (range 1.4 to 4.6), color is the discriminator pLDDT (red to blue, 40 to 90)

**FIG. S11.**
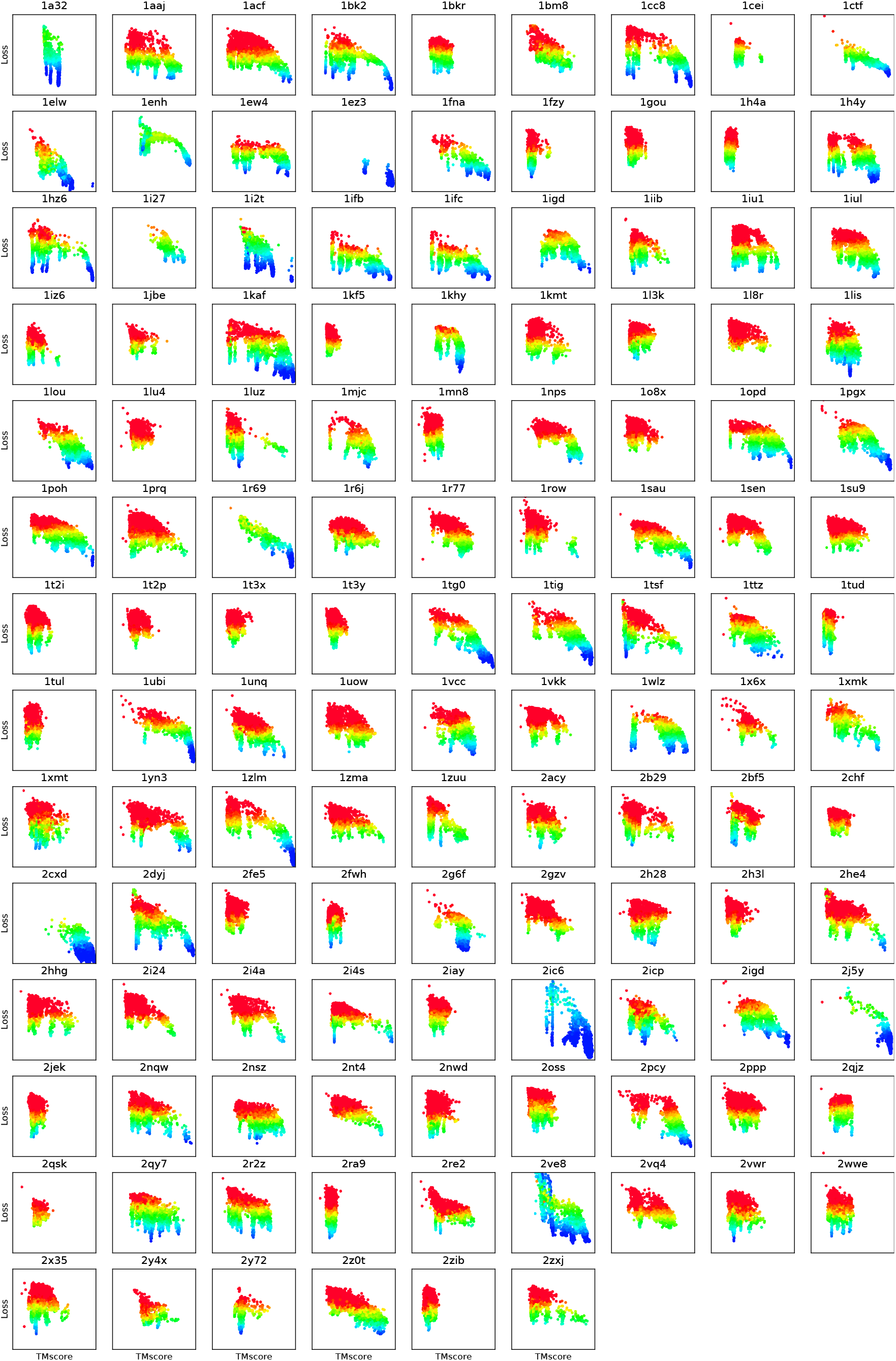
Ranking AlphaFold generated decoys with gradient descent optimization via recycling mechanism. X-axis is the TMscore (range 0 to 1), y-axis is loss (range 1.4 to 4.6), color is the discriminator pLDDT (red to blue, 40 to 90)

